# The potential of electricity transmission corridors in forested areas as bumble bee habitat

**DOI:** 10.1101/027078

**Authors:** Bruce Hill, Ignasi Bartomeus

## Abstract

Declines in pollinator abundance and diversity are not only a conservation issue but also a threat to crop pollination. Maintained infrastructure corridors, such as those containing electricity transmission lines, are potentially important wild pollinator habitat. However, there is a lack of evidence comparing the abundance and diversity of wild pollinators in transmission corridors with other important pollinator habitats. We compared the diversity of a key pollinator group, bumble bees (*Bombus spp.*), between transmission corridors and the surrounding semi-natural and managed habitat types at ten sites across Sweden’s Uppland region. Our results show that transmission corridors have no impact on bumble bee diversity in the surrounding area. However, transmission corridors and other maintained habitats have a level of bumble bees abundance and diversity comparable to semi-natural grasslands and host species that are important for conservation and ecosystem service provision. Under the current management regime, transmission corridors already provide valuable bumble bee habitat, but given that host plant density is the main determinant of bumble bee abundance, these areas could potentially be enhanced by establishing and maintaining key host plants. We show that in northern temperate regions the maintenance of transmission corridors has the potential to contribute to bumble bee conservation and the ecosystem services they provide.

## Introduction

Pollinators provide an essential ecosystem function, with 80% of plants being dependent on animal pollination for their reproduction [1]. Pollinators also provide an equally important regulating ecosystem service wherein 35% of total global crop production is reliant on animal pollination [2]. The discrepancy between supply and demand for honey bees provision of this regulating service has resulted in wild pollinators contribution to pollination gaining more recognition [3]. This is because pollination services provided by wild pollinators are often equal, complementary or superior to that provided by honey bees [4, 5]. A minority of bee species, including both managed and wild bumble bee species (*Bombus spp.*), pollinate most crops [6]. As bumble bees forage more effectively in colder temperatures than other bee species, their importance increases with latitude [7].

Pollinators are threatened by human induced environmental modification, including habitat loss, climate change and pesticides use [8, 9, 10]. Bumble bees are more sensitive to these changes than other bee species [11, 12]. Although some bumble bee species can use human modified habitats and are thriving, others are declining or near-extinct [11, 13]. For example, of the 68 bumble bee species recorded in Europe 31 species are in decline and an additional 16 species are threatened with extinction [14]. Habitat destruction [15] and a corresponding decrease of preferred host plant species [16] is one factor driving declines in bumble bee populations. For example, Europe’s semi-natural grasslands, which are a significant bumble bee nesting and foraging habitat [17, 18], have decrease by 12.8% between 1990 and 2003 [19].

In response to pollinator decline, many government and international organisations are recognising the importance of maintaining pollination services [20, 21, 22, 23]. The economic benefit provided by pollinators globally and within the EU level is estimated at €153 and €15 billion respectively [24] and therefore, maintaining and enhancing pollination is a significant area of policy. One policy response is the use of incentives. These include payments available in US through the Farm Bill 2014 [25] and in the EU through the EU Common Agricultural Policy (CAP) Agri-environmental schemes (AES). Using AES for ecological enhancement has been shown to boost bumble bee nesting and foraging habitat [26, 27, 28]. However, the potential of human-modified areas outside of agricultural land has so far received little attention from policy makers.

There is growing recognition that the routine utilitarian maintenance and disturbance of infrastructure corridors (electricity transmission corridors [29, 30, 31, 32]), roadsides [33, 34] and railway embankments [35] provides the valuable early successional landscapes required by many pollinators [36]. For example, roadside mowing has increased bee and butterfly abundance in the Netherlands [37]. Bee fauna in unmown electricity transmission corridors (hereafter transmission corridors) was richer than in adjoining annually mown grassy fields in Maryland, USA [29]. In Sweden, butterflies were more abundant in transmission corridors than in semi-natural grasslands [31, 32]. In the USA, integrated vegetation management in transmission corridors has improved the habitats of the threatened Frosted Elfin (*Callophrys irus* (Godast, 1824) and Karner Blue (*Lycaeides Melissa samuelis* (Nabokov, 1944) butterflies [38, 39].

In extensively forested parts of Europe [40, 41] and North America [38] transmission corridors can be valuable as they provide an environment suitable for herbaceous vegetation in otherwise largely forested landscapes [41]. Moreover, transmission corridors have the potential to connect discrete parts of the similar habitats [42]. However, there is limited knowledge about pollinator abundance and diversity within transmission corridors. For example, little is known about how transmission corridors compare to other pollinator habitat types and the relationship between maintenance costs of different types of infrastructure corridors and their respective pollinator abundance and diversity [36].

With the many threats to pollinators, the recognition of small-grained landscape features such as transmission corridors as valuable habitat is timely. Here, we examine the importance of transmission corridors as habitat for bumble bees, which are a key pollinator group in Sweden’s Uppland region. We compared bumble bee diversity and abundance in seven habitat types within ten spatially discrete sites - five bisected and five not bisected by transmission corridors. We predicted that transmission corridors would connect discrete patches of similar habitat and allow greater dispersal of bumble bees, consequently lowering overall beta diversity at the landscape level. However, among habitats we predicted that semi-natural grasslands would contain higher diversity compared with human-modified habitats such as transmission corridors, especially for threatened species. Finally, we reviewed the cost of maintaining and/or enhancing semi-natural grasslands and transmission corridors.

## Method and materials

### Site selection

The Swedish national transmission corridor grid (the system of 220-400 kV lines) occupies approximately 40,000 hectares, with 36,000 hectares passing through forest and consequently, requires regular maintenance. This network is owned, maintained and operated by Svenska kraftnät (SK), a state-owned public utility. SK’s transmission corridors are subject to an easement that allows them the perpetual right to construct, keep and maintain the transmission corridor grid irrespective of the underlying land tenure. In the Uppland region, transmission corridors are maintained on an eight year cycle. In year zero, transmission corridors are cleared of tall vegetation; in year three, trees threatening transmission lines are removed; in year four, transmission corridor access roads are cleared and in year seven, fast growing trees are felled. SK’s maintenance is conducted by mechanical means (J Bjermkvist, 2014 pers. comm, 3 December SK).

To investigate the influence of transmission corridors on the surrounding area, we selected ten sites of four km^2^ (2 × 2 km squares) in Sweden’s Uppland region (Supplementary Material, Figure S1). In order to minimise landscape composition confounding our results, we ensured that 1) all sites had at least 45% forest cover (range 45-70%); 2) that the second most common land use was agriculture, and 3) that all target habitats were represented (see Table 1 for habitat description). Sites were between 3.2 and 6.4 km apart. There can be a wide variation in foraging distances between bumble bee species, with radio-tracked *B. terrestris* (L, 1758) and *B. ruderatus* (Fabricius, 1775) workers foraging up to 2.5 km and 1.9 km respectively from their nests [43], while *B*. *muscorum* (L, 1758) has a much smaller foraging range of between 100-500 m [44]. Therefore, the distances between our sites minimised the chance that bumble bees recorded in one site were also recorded in another. Five sites were bisected by a transmission corridor section (widths ranging between 50-70 m), of which between 1.2-1.5 km was bordered by closed canopy forest. At the time of surveying, four sites were in year three of their maintenance schedule (all the tall vegetation was removed in 2011) and the remainder was in year six (all tall vegetation was removed in 2008). All corridors ran from north/northeast to south/southwest. The other five sites were at least 3 km from any other transmission corridors.

**Table 1:** Types of habitats and number of transects completed in each of these.

| **Transmission**<br>**corridors** | **Maintained**<br>**roadsides** | **Forest** | **Forest /**<br>**semi-**<br>**natural**<br>**grassland**<br>**boundaries** | **Semi-**<br>**natural**<br>**grasslands** | **Cereal**<br>**crop**<br>**edges** | **Maintained**<br>**drains** |
| --- | --- | --- | --- | --- | --- | --- |
| 32 | 18 | 18 | 19 | 20 | 29 | 22 |

In order to capture the variability among the surveyed habitat, we conducted multiple transects per site in each habitat (mean of 2.25 transects per habitat and site). Some sites had no representation of particular habitat types. Overall we surveyed 158 transects spread across seven habitat types (Table 1, see photos in Supplementary Material Figure S2). These habitat types were transmission corridors, semi-natural grasslands, maintained roadsides (roadsides), forest/ semi-natural grasslands boundaries, cereal crop edges, maintained drains and forests. All these habitats, except forests, have been identified as valuable bumble bee habitat in the Uppland region [17].

To our knowledge none of the surveyed transects were in areas that had been ecologically enhanced. The surveyed roadsides (all quiet tertiary or quaternary roads) are mown once annually (M. Lindqvist 2014, pers. comm, 20 May Trafikverket) whilst drains are maintained on an as-needed basis. The semi-natural grasslands surveyed met the EU’s definition of permanent pasture and grassland [45].

Each transect included an area 50 m long and up to 3m wide. All transects contained a representative density of flowering plants. Within each transect we surveyed bumble bee abundance and diversity by slowly walking along the transect for 15 minutes (a method recommended in [46]). Transects were walked twice (back and forth) but always keeping the area surveyed and the survey time fixed.

Where possible, bumble bees were identified while foraging, but most individuals could not be readily identified on the wing and therefore, were caught by net, identified and released if possible. Caught specimens that were not identified in the field were killed then identified later. Due to the difficulty distinguishing *B. terrestris* and *B. lucorum* (L. 1761) workers, all specimens were combined as *B. terrestris* [26]. Both species are common, extremely difficult to distinguish and are often grouped as they are ecologically similar. Hence, this grouping does not affect our distinction between ecosystem service providers and species of conservation concern. Collection handling time was not included in the 15 minute survey time.

When possible, the host plant of each foraging bumble bee was identified to species level during the survey, otherwise plant specimens were identified later. To correspond with peak bumble bee activity in the Uppland region [17] each site was surveyed twice between 9^th^ July 2014 and 25^th^ August 2014, with at least 2 weeks between surveys. Each survey took one day, was undertaken between 9 am and 5.30 pm and only during dry periods in temperatures above 15°C. Transects in transmission corridors were always in un-shaded areas. Before beginning each survey within the respective transect, flower density was estimated as the total percentage of the transect area covered by flowers. The categories used were “<1%”, “1–5%”, “6-10%”, “11-20%”, “21-40%”, “41-60%” and “>61%” coverage. Because all surveying were conducted by one person, this semi-quantitative measure enabled a quick yet consistent assessment of the flower density in all transects.

### Statistical analysis

To compare species abundance and richness (alpha diversity) across sites and habitats we built a generalised linear model (GLM) with species richness or abundance per transect as a function of site type (transmission corridors/no transmission corridor) and habitat type. Flower density was also included as a covariable. To account for the hierarchical structure of the data, transect nested within site was included as a random factor. Residuals were investigated to ensure they fulfilled the model assumptions and to meet the postulation of homoscedasticity we used a constant variance function. All models (see also below) were constructed using package nlme [47] in R [48]. The statistical power of the models to detect a 20% difference was calculated using package *Simr* [49].

Beta diversity was analysed on two scales. Firstly, we investigated if sites containing a transmission corridor had lower turnover rates among the different habitats. Secondly, we investigated beta diversity among different sites of the same habitat. To determine species turnover, we used additive partitioning of species richness [50, 51, 52, 53]. Alpha diversity was defined as the mean number of species per transect (i.e. species richness). The beta diversity among sites with and without transmission corridors was calculated as the total number of species found within a transmission corridor site (gamma diversity) minus the mean number of species per transect on that transmission corridor site (alpha). Beta diversity among habitats was calculated as the rarefied number of species found across all transects of a given habitat type (gamma) minus the mean number of species per transect surveyed for that habitat type (alpha). Rarefication in gamma diversity was undertaken to 90 individuals to avoid difference in sampling intensity across habitats using the package *vegan* [54] (Supplementary Material Figure S3).

From the recorded set of bumble bee species, we determined which habitats were utilised by bumble bees listed as threatened in Europe by the IUCN [14] (*B. muscorum)* and species listed as declining by Scheper et al. [16]. These included *B. humilis* (llliger, 1806), *B. sylvarum* (L, 1761) and *B. soroeensis* (Fabricius, 1777) and are hereafter termed “threatened species”. We also recorded which habitats were used by the species that are the main providers of crop pollination in Europe: *B. terrestris, B. lapidarius, B. pascuorum* (Scopoli, 1763)*, B. hypnorum* (L, 1758)*, B. pratorum* (L, 1761) and *B. hortorum* (L, 1758) [6], and are hereafter termed “provider species”. We constructed a GLM with abundance of both threatened species and provider species per transect as a function of habitat and flower density. Transect nested within site was also included as random factor. To meet the model assumptions of homoscedasticity we used a constant variance function.

Finally, to assess the importance of each host plant species for every recorded bumble bee species in the surveyed habitats, we calculated the plant species’ strengths [55] for the pool of transects of transmission corridor habitats, semi-natural grassland habitats and all habitats combined. For each plant, strength is defined as the sum of all pollinators’ dependencies on that given plant. Pollinator dependence is the fraction of all pollinator recorded visits performed on that given plant species. Therefore, a plant species could have high strength values if it attracted many pollinator species that had low dependency on it, or if it attracted few pollinators which were highly reliant on it. Note that this metric measures plant species use, not preference; a plant species could be visited by a given pollinator simply because it was the most abundant, not because it was preferred.

### Cost of managing and/or enhancing roadsides, semi-natural grasslands and transmission corridors

These managing costs were gathered from EU member material [56, 57, 58], peer-reviewed literature [26, 27, 59] and from conversations with Svenska kraftnät and Trafikverket (the Swedish Transport Administration) staff. There is large variation in the years that the management and/or enhancement costs for roadsides, semi-natural grasslands and transmission corridors were published or sourced and the initial currency in which these costs were originally stated. Therefore, no attempt was made to adjust these costs to inflation or currency fluctuations. Consequently, to enable an approximate comparison of these costs, all are expressed in Euros per hectare per annum, with the conversion of the original currency to Euros being carried out in June 2015.

## Results

In total, we recorded 1016 bumble bee specimens, comprising 20 species. These were recorded foraging on 24 plant species. Transmission corridor bisecting a site did not change bumble bee abundance (Table 2, Fig 1A) or species richness (Table 2, Fig 1B). Similarly, we found no differences among habitats in terms of total bumble bee abundance or species richness (Table 2, Fig 2A and B). As we predicted, flower density was the strongest predictor of bumble bee abundance and richness (Table 2). While the power to detect a 20% difference among sites that were bisected and not bisected by a transmission corridor is low (power ranges from 19% for abundance model to 31% for richness model), our power to detect a 20% difference between semi-natural grasslands and transmission corridors is higher (67% for the abundance model; 89% for the richness model).

**Figure 1:**
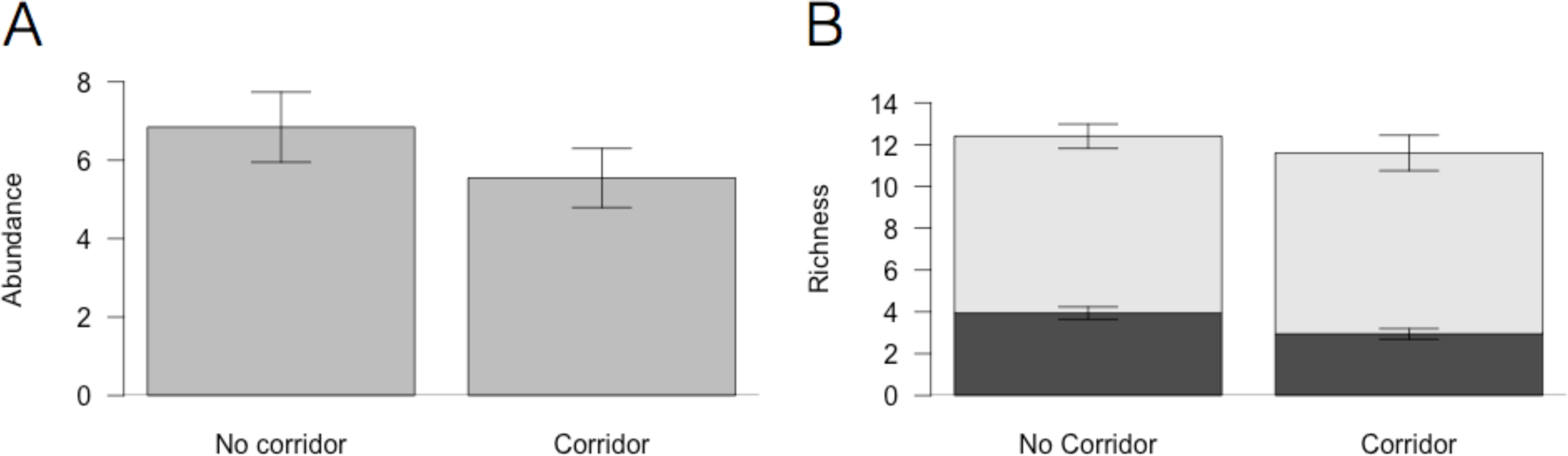
Species abundance and richness are not different in sites bisected or not bisected by a transmission corridor. A) Mean number of individuals and standard error collected per transect in transmission corridor and non transmission corridor sites. B) Mean species richness and standard error per transect in transmission corridor and non transmission corridor sites (black bars) and species beta diversity (grey bars) across habitats in sites bisected and not bisected by a transmission corridor (grey bars). The sum of both bars represents the gamma diversity of each site (n = 10 sites).

**Figure 2:**
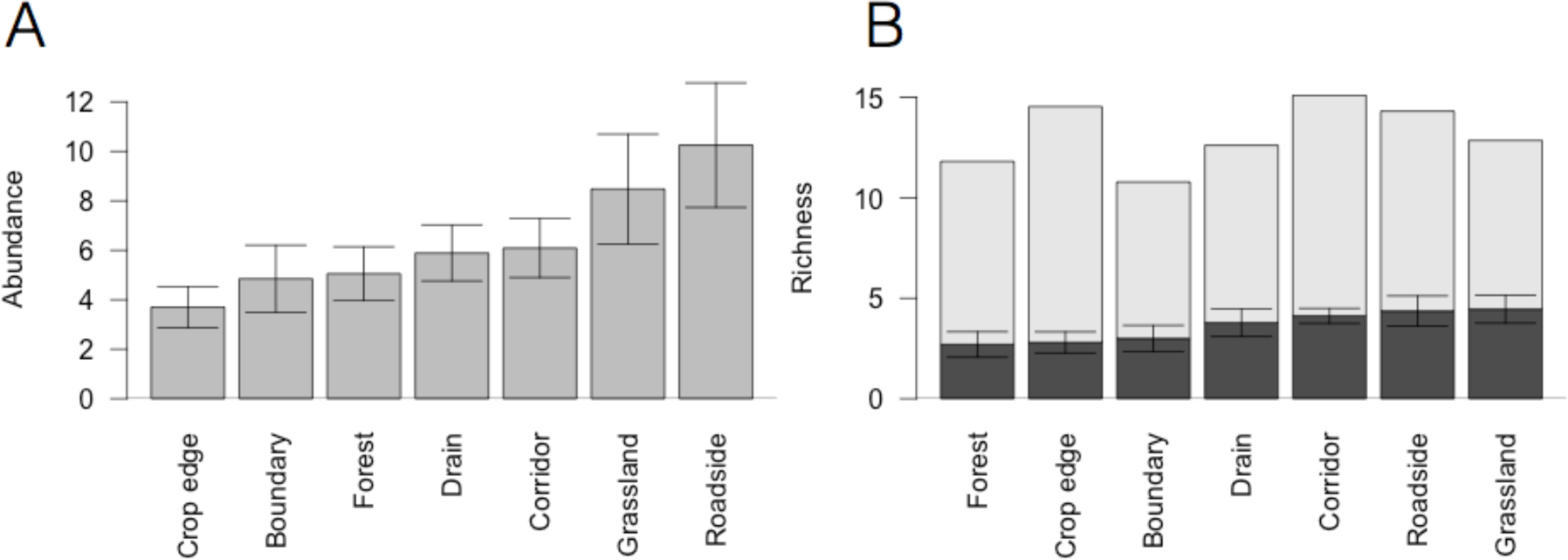
Species abundance and richness is not different across habitats. A) Mean number and standard error of individuals collected per transect in each habitat. B) Mean species richness and standard error per habitat (black bars) and species beta diversity (grey bars) between different transects of the same habitat. The sum of both bars can be seen as the gamma diversity of each habitat.

**Table 2:** Flower density is the main predictor explaining bumble bee abundances and richness. Having a transmission corridor bisecting the landscape does not increase abundance or richness. The table shows bumble bee abundance and richness models.

| **Bumble bee abundance** | **Degrees of freedom** | **F-value** | **p-value** |
| --- | --- | --- | --- |
| Flower density | 1,73 | 13.25 | <.001 |
| Habitat | 6,73 | 1.67 | 0.14 |
| Transmission corridor | 1,8 | 1.16 | 0.31 |
| **Bumble bee richness** |  |  |  |
| Flower density | 1,73 | 11.73 | 0.001 |
| Habitat | 6,73 | 1.33 | 0.25 |
| Transmission corridor | 1,8 | 2.96 | 0.12 |

Patterns of species beta diversity reveal that sites bisected by a transmission corridor did not have more homogenous species composition compared with sites not bisected by a transmission corridor (test for differences in beta diversity: n = 10, F_1,8_ = 0.03, P = 0.85, Fig 1B). We also found that species turnover among transects of the same habitat was similar, with all habitats having between 11 and 15 rarefied species (i.e. gamma diversity; Fig 2B).

We found that provider species were present in most habitats. *B. pascuorum* and *B. terrestris* were present in all habitats and were also the most abundant, while *B. lapidarius* was found in all habitats except forest. Overall, the abundance of provider species was not different across habitats (Fig 3A, Table 3). Interestingly, threatened species were not limited only to semi-natural grasslands (*B. sylvarum* and *soroeensis*), but were also found in roadsides (*B. humilis*, *soroeensis* and *sylvarum*) and transmission corridors (*B*. *muscorum* and *humilis*). However, threatened species were rarely found in the other habitat types (Fig 3B, Table 3). Flower density did not explain threatened species abundance (Table 3).

**Figure 3:**
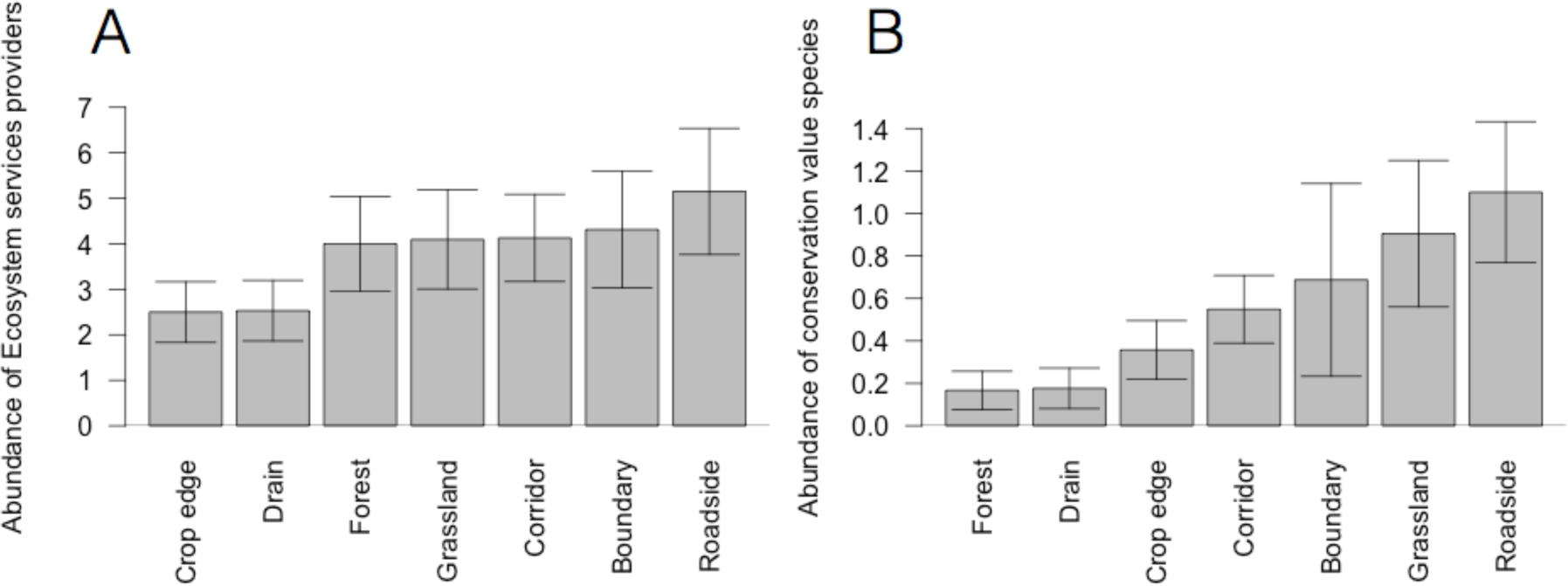
Species abundance of A) provider species is not different across habitats while for B) threatened species, transmission corridors, roadsides, semi-natural grasslands and forest-semi-natural grassland boundaries have higher abundances than the other habitats. The bars represent the mean number of individuals collected per transect in each habitat and its standard error.

**Table 3:** Abundance differences across habitats for ecosystem service providers and threatened species. While provider species mirror the general abundance pattern, for threatened species we found habitat differences, but flower cover is not longer significant.

| **Provider species abundance** | **Degrees of freedom** | **F-value** | **p-value** |
| --- | --- | --- | --- |
| Flower density | 1, 134 | 11.01 | 0.001 |
| Habitat | 6, 134 | 1.52 | 0.18 |
| **Threatened species abundance** |  |  |  |
| Flower density | 1, 62 | 0.02 | 0.89 |
| Habitat | 6, 62 | 2.72 | 0.02 |

Throughout all the sites *Carduus crispus* (L., 1753), *Trifolium pratense* (L., 1753) and *Centaurea jacea* (L., 1753) were the most important host plants for sustaining both threatened and provider species (Table 4, Fig 4). However, the importance of plant species measured as its strength varied between transmission corridors and semi-natural grasslands. For example, due to their abundance, species in the genus *Trifolium* were more important in semi-natural grasslands than in transmission corridors. Overall, important plant species sustained both bumble bee species that were not overly reliant on them and threatened species (e.g. *B. sylvarum, B. humilis*: Fig 4).

**Figure 4:**
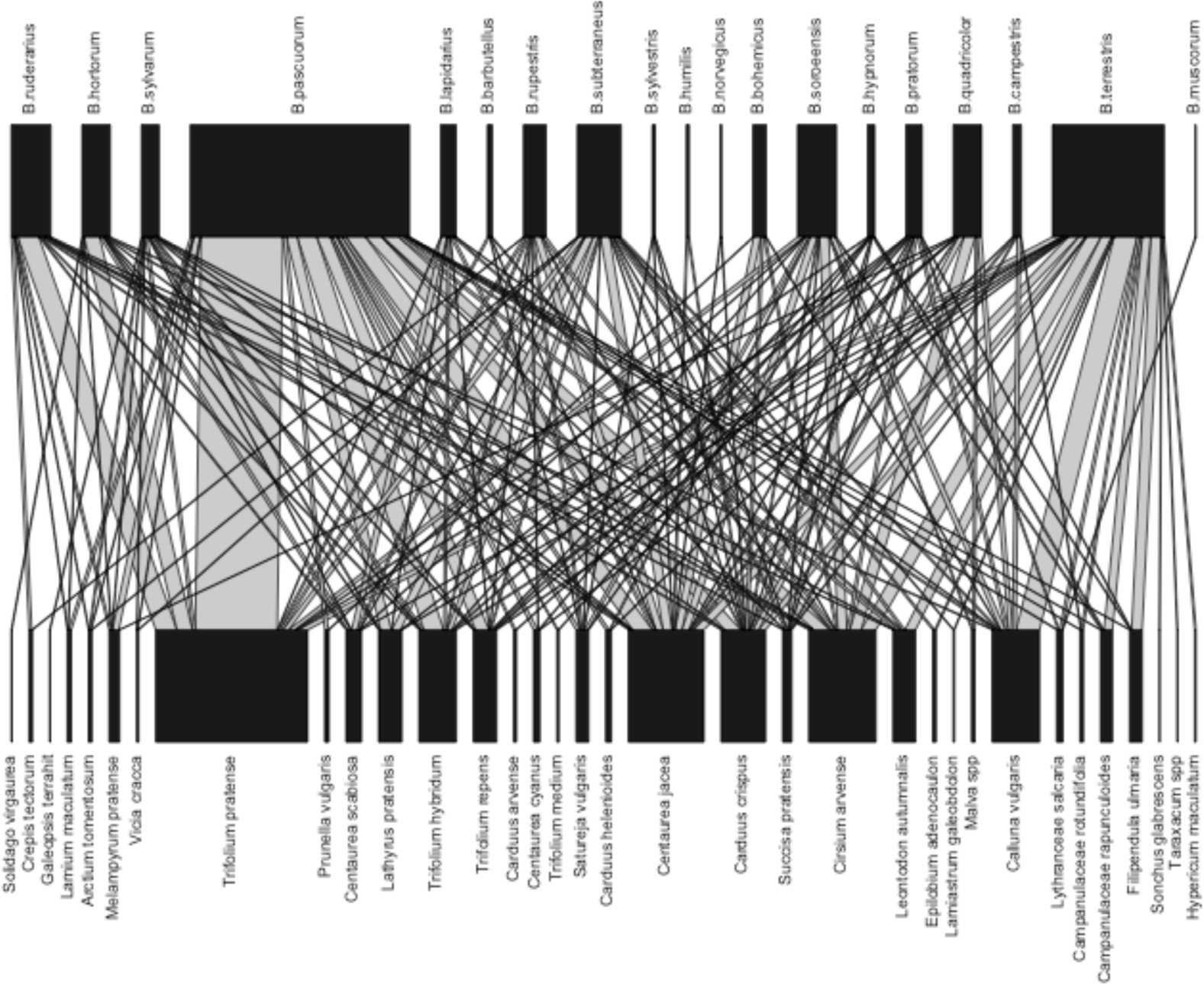
Relationship between bumble bees species and the plant species they visit. Black boxes are proportional to their total abundance. The width of the grey links between bumble bees species and the plant species they visit are proportional to the visitation frequency.

**Table 4:**
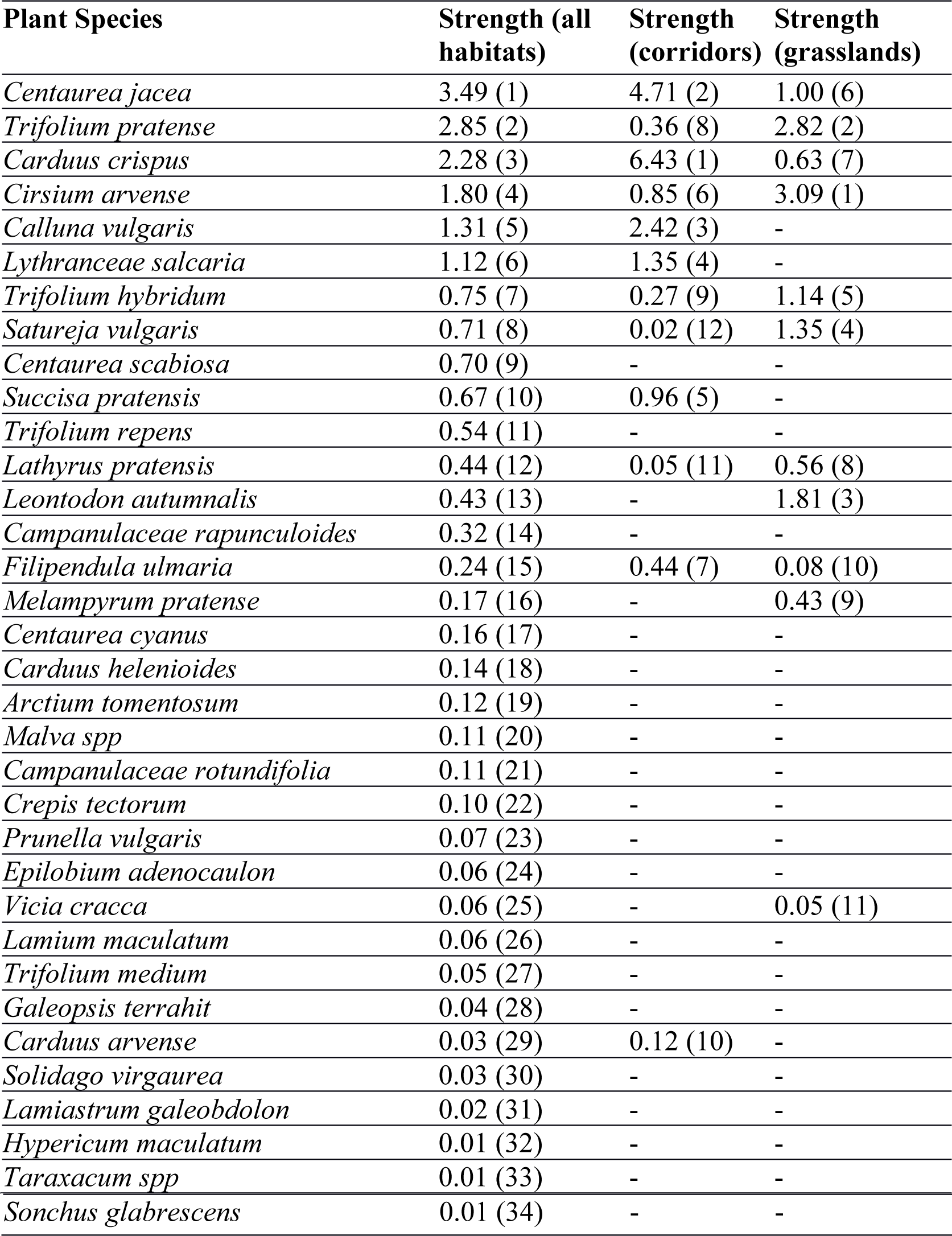
Plant species strengths (the sum of pollinator dependencies) across all interactions observed in transmission corridors, semi-natural grasslands and over all habitats combined. Rankings are in parenthesis’ because raw numbers can not be compared among habitats. Plant species with high strengths are the most important in supporting a combination of provider and threatened species. Strength values can be high because plant species support several bumble bee species with low dependence on it, or because it supports bumble bee species that are dependent on the plant species for foraging.

There was a large range in the costs of maintaining and/or ecologically enhancing transmission corridors, roadsides and semi-natural grasslands. The current maintenance of transmission corridors in Uppland costs approximately €60/ha per year (J Bjermkvist, 2014 pers comm., 3 December). Mowing Uppland roadsides similar to those surveyed costs between €500-1000/ha per year (M. Lindqvist, 2015 pers comm., 20 May). In comparison, the EU funding of Swedish AES for semi-natural grassland maintenance and enhancement, depending on inputs ranges between €121-506/ha per year [59]. Where funding is awarded, implementation of the AES is only required for 5 years [59]. This style of active management has achieved little in maintaining a diversity of grassland flora [59].

## Discussion

We found that SK’s current maintenance regime resulted in transmission corridors having bumble bee abundance and diversity equivalent to that in semi-natural grasslands. This supports the increasing recognition that transmission corridors are valuable wild pollinator habitat as in Sweden [17, 60]. In order to prevent tall vegetation damaging overhead lines, operative transmission corridors within forested areas should continue being maintained. Continuation of SK’s current management regime should result in transmission corridors providing bumble bee habitat equivalent to that supplied by semi-natural grasslands.

The fact that both transmission corridors and roadsides can sustain similar numbers of bumble bees is remarkable, especially given that the area of semi-natural grasslands in Sweden is estimated to be <10% of what it was one century ago [61]. This is particularly true for threatened species as 18 of the 41 bumble bee species in Sweden are in decline and seven more are threatened with extinction [14]. Hence, areas of transmission corridors in forested areas could provide some mitigation to the loss of semi-natural grasslands.

Roadsides also provided valuable habitat for threatened and provider species, with numbers of individuals per transect in both groups ranking higher than semi-natural grassland and forest/grassland boundaries. Roadsides tended to have high flower cover (30% density on average) which is similar to that of semi-natural grasslands. Maintained drains and cereal crop edges also had flower coverage similar to transmission corridors (13-20%), but sustained fewer bumble bee individuals, particularly those of threatened species. Dense grass swards were observed in many of the maintained drains. These swards possibly limited the habitat available for the favoured host species such as *T. pratense*, which are light demanding and low growing [62]. Overall, cereal crop edges were the narrowest habitat, with some being ≤1m wide, and hence provided the least suitable area for host plants. As forested areas of tall evergreen trees (predominantly *Pinus sylvestris* (L., 1753) and *Picea abies* (L. 1753)) had little flower cover (average of 5% density), it is not surprising that this habitat type hosted few bumble bees.

In comparison, transmission corridors and roads bisecting those forest patches were flower rich and may have an aggregation effect, concentrating pollinators into these resource rich areas [26]. However, it is important to note that flower density did not explain threatened species abundance, which suggests other factors, such as nesting sites, may be more limiting for these species [26]. It is not known what the effects of electrical and magnetic field radiation from high voltage powerlines have on bees [36] and quiet roads potentially represent a minor threat to bumble bees [33]. It is possible that these risks are countered in transmission corridors by providing suitable habitat for rodents, thereby potentially increasing nesting availability for bumble bees using abandoned rodent cavities as nesting sites [63]. Similarly, roadsides often contain areas of withered grass and tussocks that are crucial for nesting sites [17].

Overall, our results do not indicate that transmission corridors enhance bumble bee abundance or species richness by increasing connectivity of non-forested habitats or by having a spill-over effect into surrounding habitats. However, with only 10 sites the power to detect such landscape effects in our dataset is limited. The intrinsic variability in bumble bee populations between years [64] suggests that long term data in different boreal countries are needed to confirm our results.

Within transmission corridors the main host plants for bumble bees are mostly limited to small areas that are not dominated by shading shrubby vegetation (B. Hill 2015 pers. obs, 12 November). Floral density is an important predictor of bumble bee diversity and abundance. The large areas of herbaceous vegetation and shrubs within transmission corridors could provide considerable potential to enhance bumble bee habitat. Such actions could also assist in providing the ~2% of flower-rich habitat within farmland that is required to maintain provider bumble bee species colonies [65].

Maintaining and enhancing the abundance of early flowering *Salix* species such as *Salix caprea* (L. 1753) is a way of potentially improving the quality of bumble habitat in transmission corridors. Early flowering *Salix* species provide critical forage for early emerging bumble bee queens and subsequently, successful colony establishment. It has been shown that >1000m^3^ crown volume/ha positively influenced bumble bee abundance [17]. Flower abundance later in the season is also critical for late emerging species because many of these are threatened [16]. In Sweden, bumble bees are mostly active up to early September, after which the new queens hibernate underground [17]. As we surveyed almost to this period, we assume that we captured the peak phenology of most bumble bee species, including the threatened species.

For most bumble bee species, legumes and other nectar rich flowers are a significant resource [62] and our results support this observation. Although we did not separate nectar and pollen foraging trips, it is likely that different plant species are important for different reasons. For example, while *T. pratense* is a rich source of nectar and pollen, most thistle species may be used only for nectar [62]. However, in comparison to semi-natural grasslands, the transmission corridors we surveyed had a lower abundance of key plants such as *T. pratense*. The sowing of nectar rich flower seeds is a proven way of enhancing bumble bee abundance and diversity [28]. This is a possible means of enhancing bumble bee habitat in transmission corridors and would cost approximately €42/ha/yr [58]. Suitable open areas include access roads as these are not dominated by shading shrubby vegetation, and the additional areas of bare earth exposed during their maintenance.

Increasing the amount of open habitat within transmission corridors is another potential way of increasing host plant habitat and consequently, bumble bee diversity and abundance [29, 37, 65]. Removal of existing shrubs on transmission corridors would cost approximately €14/ha/yr [66]. Host plants might then naturally colonise these areas or seeds of suitable species could be sown.

Funding the enhancement of bumble bee habitat within transmission corridors could be an effective way to both benefit bumble bee conservation and increase the pollination services they provide. It might also augment the ecological value of these areas. Depending on the location, enhancing the ecological value of transmission corridors could be conducted in tandem with the protection of ecological focus areas as prescribed by the EU [45]. The opportunity cost of producing an ecological focus area via converting productive agricultural land to unproductive biodiversity rich areas can be considerable. For example, winter wheat which is a major crop in Uppland region, can provide gross returns of between €565/ha-€1505/ha [67, 68]. The establishment and maintenance of biodiversity rich areas within transmission corridors, like those studied here, would avoid any such opportunity cost. The permanence of transmission corridors in the landscape also means that any enhancement within these is likely to provide long-term benefits. Such actions might well aid in meeting the EU’s *Biodiversity Strategy to 2020* Target 2, as well as the 2020 headline target [20]. However, areas of transmission corridors do not meet the EU’s CAP, enabling definitions of either “eligible hectare” or “ecological focus area”. Therefore, funding via EU AES for the ecological enhancement of such areas is not currently possible [45].

Pollinator habitat within transmission corridors is spatially limited to certain areas. Moreover, we only tested for the effect of transmission corridors in forested landscapes. The ability of transmission corridors to sustain pollinators in non-forested landscapes is still unexplored. Consequently, transmission corridors cannot substitute AES, but can complement it. In other situations it has been shown that tailoring inputs for specific results is possible. Application of AES is simple - resource poor landscapes e.g. croplands had the greatest benefit to provider species, whilst applying AES in more complex landscapes provided more benefit to threatened species [69]. The widespread geographic extent of transmission corridors through many northern hemisphere landscapes provides valuable but yet to be fully exploited opportunities for bumble bee conservation. However the benefit of transmission corridors for biodiversity other than bumble bees has not yet been explored.

## Conclusions

Bumble bee abundance and diversity is threatened by many factors. Given both the intrinsic value of bumble bees and the ecosystem service they provide, actions are being taken to counter these threats. Studies, including ours have shown that the maintenance of transmission and other infrastructure corridors may unintentionally create valuable habitat for pollinators. Our study also shows that SK’s current transmission corridor maintenance regime is a cost effective way of producing such habitat when compared to other maintenance regimes. The permanence and extent of transmission corridors means that any wild pollinator habitat created due to their maintenance is likely to be present long-term. There are simple, proven management practices to enhance bumble bee richness and abundance but further research is needed to evaluate and optimise conservation approaches. Funding is needed for such work. Any future reviews of the Europe 2020 Strategy, CAP, or similar policy may provide opportunities to promote incentives to enhance the valuable pollinator habitat provided by maintaining infrastructure corridors.

## Data, code and materials

All data and code to reproduce this analysis are deposited in <u>www.github.com/ibartomeus/powerlines</u>. Data available from the Dryad Digital Repository: http://dx.doi.org/10.5061/dryad.v32df

## Competing interests

We have no competing interests

## Author contributions

BH and IB conceived the study; BH collected field data, participated in the design of the study and drafted the manuscript; IB designed the study, carried out the statistical analyses, coordinated the study and helped draft the manuscript. All authors gave final approval for publication.

## Acknowledgements

We thank WSP Sverige for providing logistic support. We thank David Kleijn and two anonymous reviewers for comments in a previous draft, Jamie Stavert for English language editing and Gerald Malsher and Björn Cederberg (Sveriges lantbruksuniversitet/Swedish University of Agricultural Sciences) for identifying several bumble bee specimens.

## Funding

BH was funded by Svenska kraftnät solely for transport and field expenses. IB was funded by EU project BeeFun (PCIG14-GA-2013-631653). Svenska kraftnät took no part in experimental design or data interpretation.

